# iPSCs-derived Bronchial Airways-On-Chip for Assessment of Cytokine Secretion triggered by Volatile Organic Compounds

**DOI:** 10.1101/2025.09.07.674694

**Authors:** Adele Goldman-Pinkovich, Rose Ibraheem-Azaizeh, Yasmin Habib, Abeer Abassi, Shir Shapiro-Fillin, Ronit Almog, Josué Sznitman, Arbel Artzy-Schnirman

## Abstract

Air quality monitoring currently relies mostly on a combination of epidemiological data and classic experimental data. Our objective was to design an alternative approach for assessing air pollutant risk potential using a specialized platform capable of detecting long-term and indirect effects of exposure via the inhaled route. We used a Bronchial Airways-On-Chip (BOC) offering some of the physiological complexity of the human lung based on our previously developed device coupled with in-vitro differentiated bronchial epithelium derived from induced Pluripotent Stem Cells (iPSCs). This setup is capable of replicating bronchial epithelia exposure to irritants at air-liquid interface, in controlled and reproducible conditions. ***It comprises the first proof-of-concept design combining a BOC with iPSCs-derived bronchial epithelium as an alternative approach towards potential risk assessmnet of inhaled pollutants***. As a representative pollutant we use Benzene, a Volatile Organic Compound (VOC). At low concentrations and short-term exposure it is not considered acutely harmful, but long-term exposure can result in mutagenic and carcinogenic effects. As air pollutant toxicity is known to be mediated by the respiratory epithelial lining and secretion of cytokines, we demonstrate our system to be sufficiently sensitive to capture increased cytokine secretion corresponding to increasing concentrations of Benzene. ***Of utmost relevance is our finding that an accumulative effect could be detected, only caused by prolonged exposure at low concentrations of Benzene, previously shown to be non-toxic in classic short-term in-vitro studies***. Finally, the accumulative effect could be reversed using a commonly-used asthma medication (Montelukast), further supporting the relevance of the setup.

## 2. INTRODUCTION

Smoking and air pollution rank as the fourth leading cause of death globally, accounting for approximately 5% of annual deaths worldwide^1^. According to the World Health Organization (WHO), air pollution is the largest environmental health risk and a major contributor to premature morbidity and mortality^2^. Air pollution consists of a complex mixture of gases and particulate matter, including various chemical compounds that pose significant risks to respiratory health. Among these, Volatile Organic Compounds (VOCs) such as benzene are of particular concern due to their widespread presence in cigarette smoke, vehicle exhaust, and industrial emissions. VOCs contribute substantially to pulmonary and cardiovascular diseases, and long-term exposure even at low concentrations can lead to mutagenic and carcinogenic effects^3^.

Accurate air quality monitoring and establishing thresholds for acceptable pollutant levels are therefore critical. Current assessments rely on studies of inhaled pollutant toxicity and exposure patterns using a combination of classical cell cultures, animal models, and epidemiological data^2,4,5^. However, these methods have limitations: on the one hand, static *in vitro* cell cultures fail to recapitulate the structural and functional complexity of the human lung^6,7^, whereas animal models often lack direct translational relevance to human physiology^8,9^. Finally, epidemiological studies in humans often have difficulties with variable isolation and establishing causality^10^.

To address these gaps, Organ-On-Chip (OOC) platforms have emerged as promising *in vitro* tools that improve physiological relevance, standardization, and experimental control^11,12^ while mimicking closely human cellular makeup. In particular, Airways-On-Chip (AOC) systems offer unique advantages over traditional models by enabling cellular exposure to pollutants via inhalation at an air-liquid interface (ALI)^13^. Unlike conventional cultures, AOCs can replicate physiological inhalation airflows at true scale, a key factor influencing particle deposition on the airway lumen^14^, that also influence biological features such as mucin secretion, ciliated cell differentiation, and epithelial glycocalyx formation^15–17^.

Several groups, including our own, have developed AOCs using primary bronchial epithelial cells. These biomimetic lung-on-chip devices have been applied to model pulmonary diseases such as cystic fibrosis (CF)^18^ and primary ciliary dyskinesia (PCD)^19^, as well as to study tobacco smoke and air pollution hazards^20^. Notably, Benam et al. (2016) demonstrated chronic obstructive pulmonary disease (COPD) modeling by culturing primary cells from COPD patients in microfluidic devices under ALI conditions, combined with whole cigarette smoke exposure using a micro-respirator. These microphysiological lung platforms allow direct comparisons of epithelial responses from the same donor with and without smoke exposure, revealing disease-specific molecular signatures and inflammatory pathways^20^. The use of primary bronchial epithelial cells from patient biopsies, however, encompasses some limitations such as donor availability, variability, and passage number. Induced Pluripotent Stem Cells (iPSCs)^36^ for *in-vitro* differentiation of bronchial epithelium have been shown to provide the major components (see ***Fig. 1b, Fig. 2b***). of the pseudostratified bronchial epithelium as it occurs naturally while offering significant advantages such as incomparable accessibility and standardization. We utilize this approach in our models and verify normal epithelial layer composition to support its physiological relevance.

**Figure 1.**
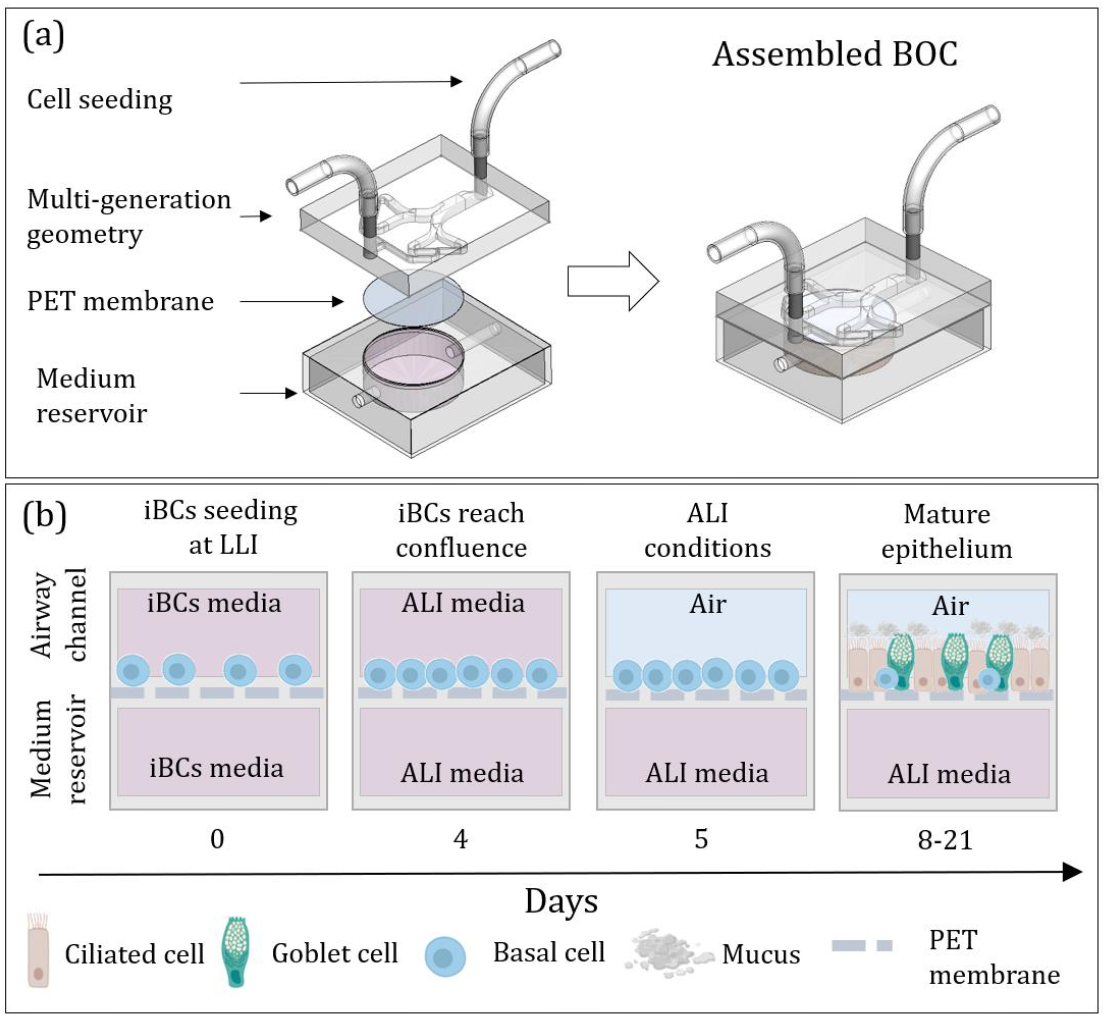
Bronchial Airway-On-Chip (BOC) model of the upper small bronchial airway. **(a)** Components of the sandwich structure of the model and collapsed assembly. **(b)** Stages of iBCs-derived epithelial maturation at ALI initiated directly in the model by seeding iBCs in the airway channel. Based on Suzuki et al., 2021.

**Figure 2.**
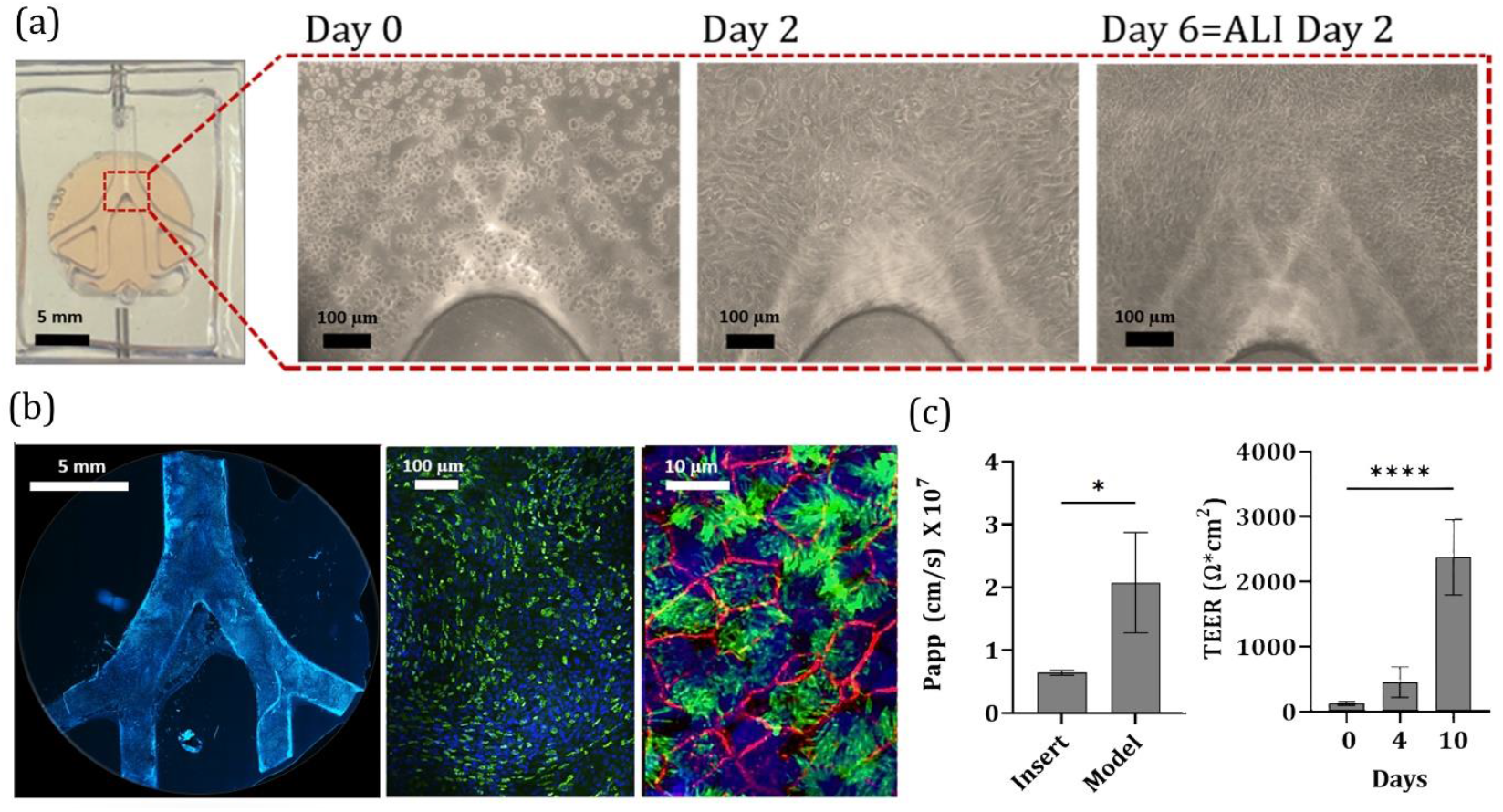
Bronchial Airway-On-Chip model of the small airway. **(a)** iBCs-derived epithelial maturation at ALI within the model. Model seeded with iBCs and epithelial differentiation process shown from seeding to maturation in bright field. Scale bars indicate 100 µm. **(b)** Left to right: DAPI stained nuclei of cells (left panel, blue) confluently seeded throughout the Y-shape, imaged using the X10 objective to show delicate ciliary structures (middle panel, green). An X63 objective was used to further demonstrate cilia (green) and tight junctions (right panel, red). **(c**) **Right panel**: Apparent permeability (*P*_app_ analysis using fluorescein) comparison of mature epithelia differentiated between models and inserts. Unpaired t-test was significant (n=3, *p-value<0.05). **Left panel:**. TEER was measured routinely in trans-wells prior to transferring the cells to ALI conditions and on days 8-21 to verify normal epithelial layer integrity and healthy maturation progress. A TEER of >400 Ω*cm^2^ on days 3-4 after seeding is considered as a threshold for normal iBCs attachment and proliferation prior to proceeding to ALI conditions and completing the epithelial maturation process. Unpaired t-test was significant (n=5, ****p-value<0.0001).

The airway epithelium forms the first barrier against inhaled irritants and pathogens and plays a critical role in the innate immune response^21^. Upon exposure, epithelial cells secrete cytokines that mediate the onset of inflammatory cascades, making cytokine profiles valuable indicators of tissue response and inflammation severity^22–25^. Lung-on-chip platforms frequently assess cytokine secretion to evaluate pollutant and cigarette smoke effects in both healthy and diseased contexts such as asthma and COPD^20^. Specifically, cytokines involved in the bronchial response to pollution and related diseases include e.g. IL-8 and IL-1β as pro-inflammatory markers, and IL-10, which exhibits a complex role as both an anti-inflammatory and pro-inflammatory cytokine depending on context^26,27^. Both downregulation and over-stimulation of IL-10 secretion can have both a protective and a harmful effect^28^. In the lung there is evidence to support involvement of IL-10 in mucus metaplasia, tissue inflammation, and airway remodeling^29^ as well as Neutrophil Extracellular Traps (NETs) formation^30^. IL-10 suppresses the Reactive Oxygen Species (ROS)-dependent formation of NETs whereas IL-10 downregulation is linked to exacerbated lung fibrosis and cystic fibrosis^31^.

Motivated by ongoing respiratory health concerns and ensuing pulmonary diseases resulting from environmental exposure, in the present study we leverage the potential of an induced pluripotent stem cell (iPSC)-derived bronchial AOC model to assess subtle inflammatory effects of airborne VOC exposure. Specifically we concentrate on benzene, a representative VOC recognized as a human carcinogen commonly found in cigarette smoke, vehicle exhaust, and industrial emissions^32,33^. While at low concentrations some VOCs are not considered acutely harmful to human health with short-term exposure, long-term exposure may result in mutagenic and carcinogenic effects^34^. The WHO considers benzene exposure above1.7 µg/L (<10^−8^ M) associated with increased lifetime cancer risk by 1/100,000^2^. In past *in vitro* studies this level has not been considered directly cytotoxic^4^. Using our airway-on-chip platform integrating iPSC-derived bronchial epithelial cells, we investigate how sub-cytotoxic benzene exposure under inhalation airflows affects cytokine secretion (IL-1β, IL-8, and IL-10). By isolating early epithelial-pollutant interactions, our *in vitro* platform enables mechanistic insights into inflammation initiation and the pathways leading to long-term pulmonary pathology. This approach complements existing toxicity assessment methods and advances the development of human-relevant models for environmental health risk evaluation.

## 3. RESULTS

### 2.2.1 Bronchial Epithelium Maturation in a Bronchial Airway-On-Chip

Building on previous work in the group, we leveraged a previously developed bronchial *airway-on-chip* featuring two planar, symmetric branching airway channels, resembling the anatomy of small bronchial airways allowing for airflow within the model^37,38^ (***Figure 1a, Table S1***). Cells were seeded in the model by injecting induced basal cells (iBCs) in the Y-channel and allowing for attachment and confluency before transitioning to ALI (***Figure 1b***).

As part of the calibration process, we characterized the mature epithelial layer in terms of ciliated and goblet cell percentage (***Table 1, Figure 2***) as well as marker expression (***Figure 3a and 3b***) and layer integrity (***Figure 3c***). Vortex-shaped muco-ciliary patterning is expected, indicative of the cilia–mucus hydrodynamic interactions resulting from the presence of a ciliary-beat^17^ (***Figure S1e***). We found however that this feature is somewhat affected by the geometry of the containing vessel (i.e. airway channel versus trans-well) so the classically round vortex shaped patterning was more prominent and could be captured more clearly in trans-wells. Layer integrity was determined by measuring apparent permeability (*P*_*aap*_) and Trans-epithelial Electrical Resistance (TEER). TEER is measured only in trans-wells due to technical limitations (***Table 1, Figure 2, Figure S1***). Indeed, layer integrity by *P*_app_ (***Figure 2e***) exhibited only a minor difference (of`2.63-fold increase) compared to inserts. Layer composition was also unaffected (***Table 1, Figure 2 and Figure S1***). Epithelial markers expression was validated both by RT-PCR (in inserts; ***Figure 3***) and Immunohistochemistry (***Figure 2c, Supplementary Figure S1***).

**Table 1.** Epithelial Layer Composition and CBF.

| Marker |  | Human Bronchus<br>(small airway epithelium) | iBCs-derived EP |
| --- | --- | --- | --- |
| Ciliated Cells | α-tubulin (%) | 73 (1)<sup>39</sup> | 66 (18) |
| Goblet Cells | MUC5AC (%) | 10.3<sup>40</sup> | 13.1 (5.7)* |
| Layer Integrity | $P_{app}$ (cm/s) $\times 10^7$ | - | 2.1 (0.8) |
| | TEER ($\Omega \cdot \text{cm}^2$) | 700-1200<sup>41</sup> | 2375.4 (580.41)* |
| CBF | beats/sec | 9-20<sup>42</sup> | 16 (3)* |

\* Values were determined in trans-wells

**Figure 3.**
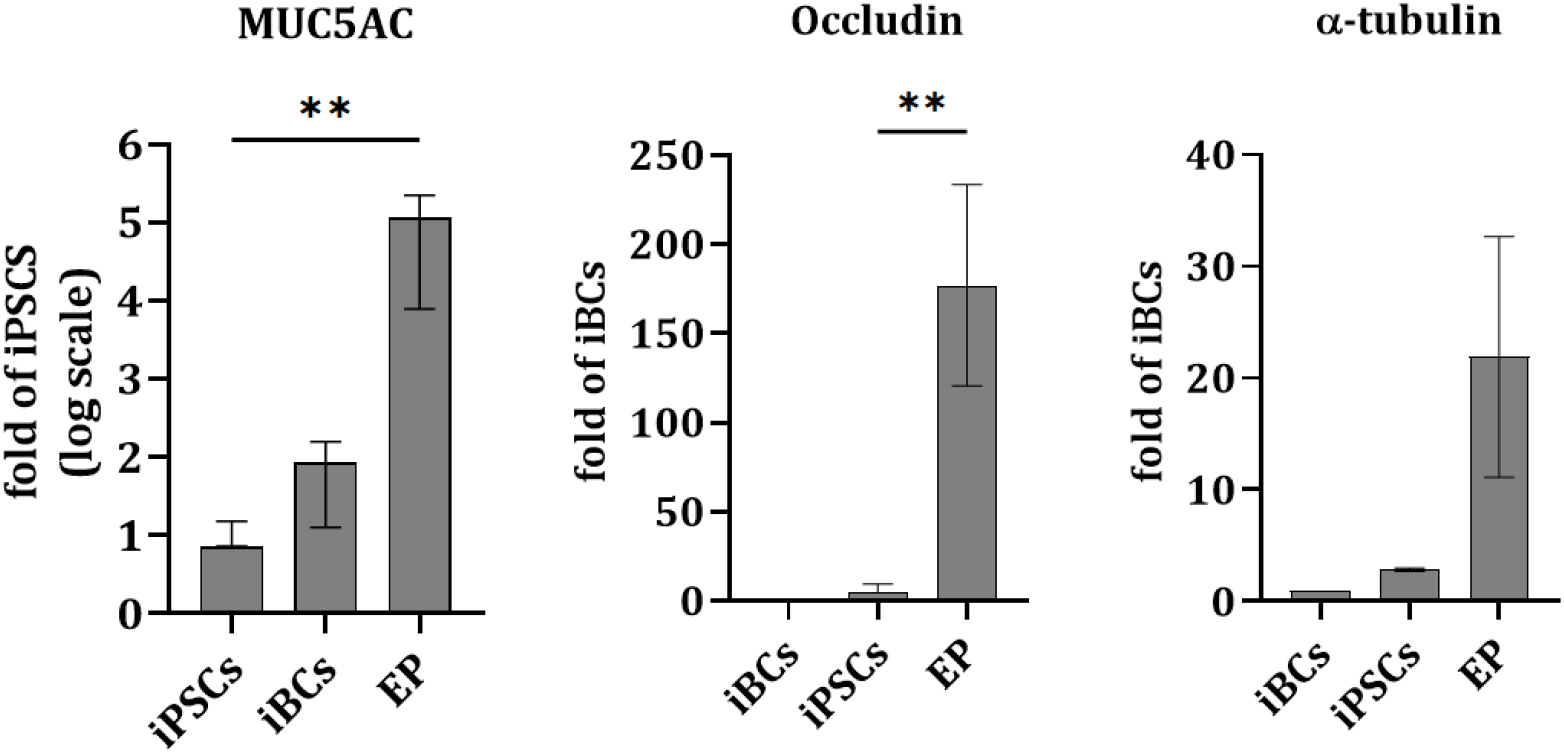
Stage specific markers expression validation. Epithelial marker expression in mature airway epithelium (EP) was compared to that of undifferentiated iPSCs and iBCs (n=3) using RT-PCR. Unpaired t-test was significant (n=3, **p-value<0.01).

### 2.2.2 From iPSCs to iBCs

To the best of our knowledge the vast majority of existing research utilizing bronchial epithelium and lung-on-chip platforms use either primary cells or cell lines, apart from the work of Yadav et al., 2025^43^ who have used in-vitro differentiated bronchial epithelia derived from iPSCs. This approach holds a vast array of advantages. The accessibility of differentiating mature epithelium from a cryopreserved stock of airway progenitors, i.e., iBCs that are not immortalized cells is unprecedented. Furthermore, it eliminates donor variability and improves experimental reproducibility and standardization. Finally, it paves the way towards patient-specific experimental set up, and as iPSCs can be derived from peripheral blood it also eliminates the need for invasive biopsies.

Here we differentiated mature bronchial epithelium from iPSCs derived from healthy donor skin fibroblasts^44^ as per the protocol of Suzuki et al., 2021^45^, and the relevant milestones were tested as depicted in detail in ***Figures 1 and 3, Figures S1 and S2*** and in methods section 2.1.2. Basal cells are naturally occurring airway progenitors that line the bronchial airways and regularly differentiate to give rise to mature bronchial epithelia. iBCs are selected following a timely protocol of exposure to signal molecules and subsequent isolation of cells that are positive for several markers such as NKX2.1/TTF1 transcription factor, TP63 (Tumor Protein P63), NGFR (Nerve Growth Factor Receptor), EpCAM (Epithelial Cell Adhesion Molecule) and CPM (Carboxypeptidase M). The latter three are membranal proteins that are therefore used for live cell selection (***Figure S1***). In this setup we substituted FACS-sorting for magnetic beads sorting is noteworthy; this improved time and cost efficiency while maintaining efficiency as shown in the comparison of the two methods (***Figure S2***). All data presented herein has been collected using magnetic beads enrichment both during differentiation and routing maintenance.

### 2.2.3 Cytokine secretion upon Benzene exposure

Cytokine secretion was determined at two concentrations of benzene that were previously shown to be non-toxic (10^-8^ M) or toxic (10^-4^ M) in a short-term exposure set-up^4^. Our data indicated that IL-10 variability from control levels (no benzene exposure) could be noted even at the non-toxic concentration of 10^-6^ M, but not at 10^-8^ M (***Figure 4a; Supplementary Figure S4***). However, in the case of inflammatory cytokines IL-1β and IL-8 no significant effect was found at any of the concentrations examined.

**Figure 4.**
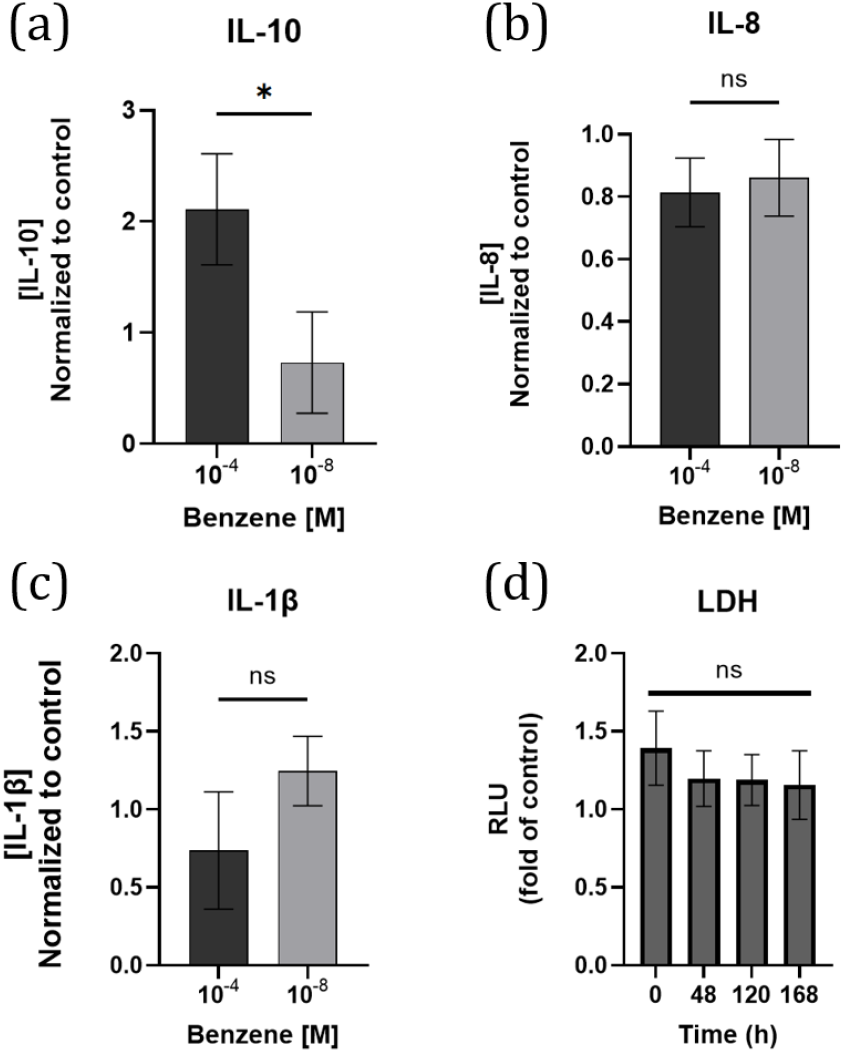
Dose-dependent cytokine secretion following benzene exposure. Benzene at the indicated concentrations or media only (control) was introduced into the airway channel for 1 hour. ELISA (a, b, c) and LDH Cytotoxicity (d) assays were carried out 48 hours post exposure (or as indicated) to measure IL-10 (a) IL-1β (b) or IL-8 (c) secretion into the basal chamber media and assess cell viability. (d) Viability RFU values are relative to paired models at each time point treated with non-toxic benzene concentration (10^-8^ M). For all time points the minimal number of different biological repeats (models) was n≥3. One-way ANOVA: ns.

A major advantage of the use of the present models is that they allow for a physiologically-relevant route of exposure that can simulate these cumulative effects of exposure to pollutants. We were therefore interested in demonstrating an accumulative effect of exposure to benzene, and introduced additional benzene exposures at 48, 72 and 120 hours after the initial exposure (***Figure 5***). Indeed, in this case we found increased IL-10 secretion even at the concentration of 10^-8^M, a non-toxic concentration that did not lead to higher levels of IL-10 secretion versus those of control when tested only at 48 hours after a single benzene exposure (***Figure 4a***). The addition of Montelukast at 120 hours following repeated exposure to 10^-4^M benzene has reduced IL-10 levels at 168 significantly (***Figure 5b)***.

**Figure 5.**
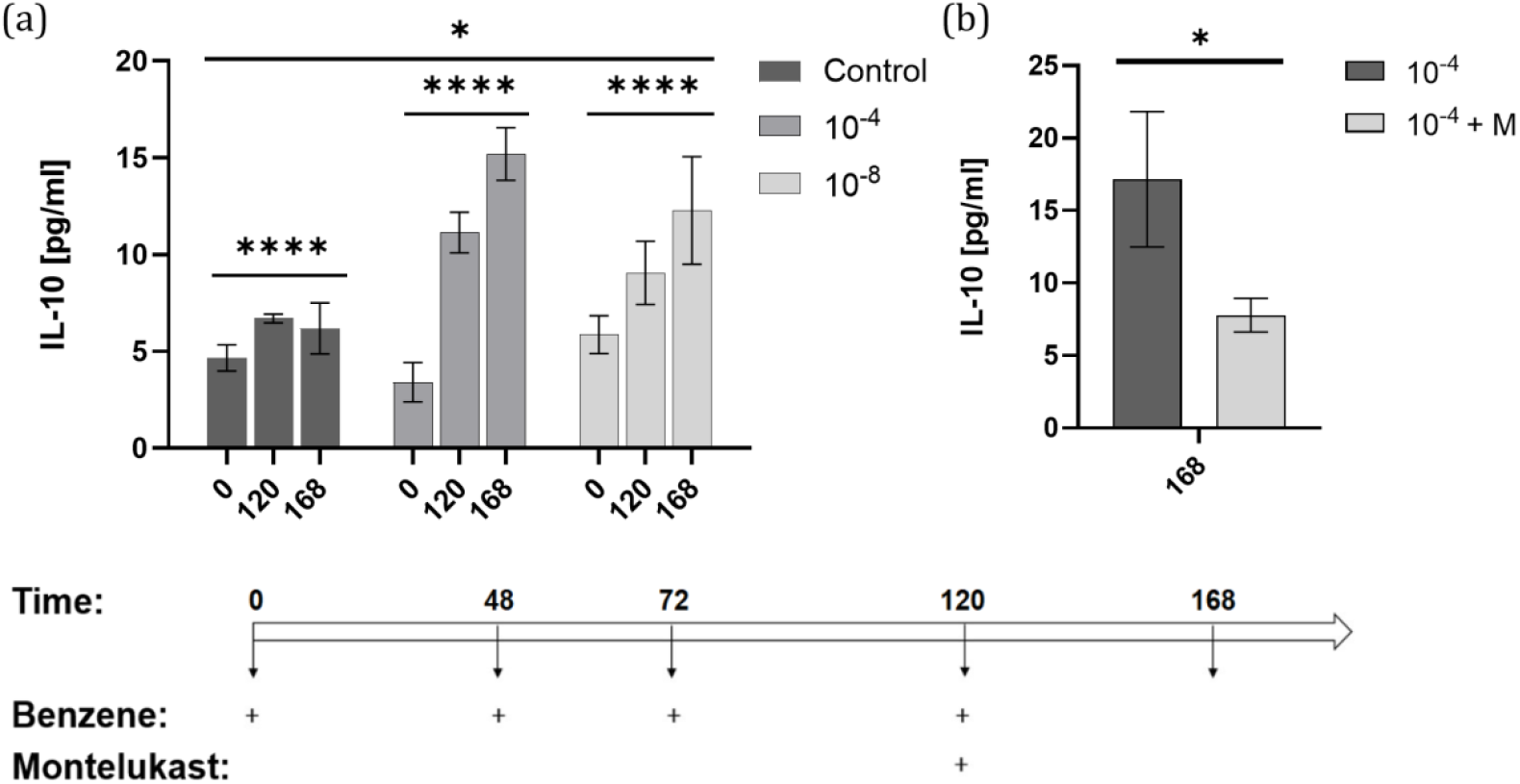
Cumulative effect of benzene exposure. (a) Repeated exposure to benzene at ALI media at concentrations of 10^-4^ M or 10^-8^ M via the airway channel resulted in an accumulative effect. Exposure was carried out at the indicated several time points post initial exposure (48, 72 and 120 hours) and cytokine secretions was determined in media from the basal compartment at 120 and 168 hours following initial exposure. Number of biological repeats for each condition is n≥3. Two-way ANOVA analysis was significant both for the effect of time of exposure (****p-value<0.0001) and Benzene concentration (*p-value=0.005). Interaction (p-value<0.05). (b) IL-10 secretion following exposure to Benzene at 10^-4^ M with or without the addition of Montelukast (one-tailed ns *p-value=0.058).

## DISCUSSION

The airway epithelium forms a robust barrier protecting the body from inhaled toxic particles and pathogens. Air pollutant toxicity is known to be mediated by the respiratory epithelial lining for recruitment of immune components and consequent secretion of cytokines. IL-10 is often regarded as an anti-inflammatory cytokine but has been suggested to play a dual-role in inflammatory response^26,27^. Both downregulation and over-stimulation of IL-10 secretion can have both a protective and a harmful effect^28^. In the lung there is evidence to support involvement of IL-10 in mucus metaplasia, tissue inflammation, and airway remodeling^29^ as well as Neutrophil Extracellular Traps (NETs) formation^30^. IL-10 suppresses the Reactive Oxygen Species (ROS)-dependent formation of NETs whereas IL-10 downregulation is linked to exacerbated lung fibrosis and cystic fibrosis^31^. IL-10, which has been shown to play both protective and pathogenic roles in lung inflammation, airway remodeling, and immune suppression is often considered as the most potent anti-inflammatory cytokine and is recognized for enabling cancer immune surveillance and tumor rejection ^46^. As a proof-of-concept we used Benzene as a representative VOC to study air pollution exposure in our platform since it is a common air pollutant with known carcinogenic potential, yet it is not considered acutely toxic. It has previously been assessed for toxicity in epithelial cultures and its accumulative affects by exposure via the inhaled route as well as by skin contact have been characterized^4,47^. While there is evidence to suggest benzene exposure of airway epithelia (such as through cigarette smoke) induces pro-inflammatory cytokine release (IL-1b^48–50^, IL-8^48,50^) others suggest that sub-cytotoxic benzene exposure can cause disruptions (such as oxidative stress signaling and immune cell chemotaxis) in airway epithelial cells without activating typical inflammatory of metabolic pathways^51^. IL-10 has a sensitive and regulatory role in allergic airway inflammation and benzene exposure. Studies show that IL-10 is elevated in sera of Benzene exposed individuals and that injecting anti-IL-10 antibodies into benzene-treated mice can reverse some of the immune suppression inflicted by Benzene exposure^50,52^. However in asthma or COPD, conditions where benzene exposure has been shown to play a role as part of general air pollution, IL-10 is generally found to be downregulated^53^.

Furtheremore, benzene exposure leading towards immune suppression can be reversed by IL-10 removal^50^. This is in line with our data showing no signifact response of pro-inflammatory cytokine secretion in response to benzene exposure but a clear IL-10 response was evident (***Figure 4***). It is noteworthy to recall that our system only includes the innate component of the bronchial epithelial barrier involved in the first encounter with benzene. This way we are able to address the very initial steps of an effect that is likely further complicated and effected by downstream components of the full complexity of an organism. No existing model can offer a realistic way to test an accumulating effect of persistent exposure to very small amounts of pollutants, which occurs in densely populated and commercial zones. Our data suggest that using *in vitro* models such as ours can help overcome this gap and pinpoint cumulative effects, which might be masked in animal models or epidemiological investigations due to their complexity.

In the context of airway diseases, it has been shown that treatment with Montelukast (an antagonist of cysteinyl leukotrienes receptor 1) in asthmatic children improves their impaired immunological balance through the increase of serum IL-10^54,55^. The opposite effect can also be observed as in the case of LPS-induced IL-10 expression that was suppressed by montelukast in M2 macrophages *in vitro*^56^. One may resolve this apparent contradiction by examining the nature of the baseline state of each system. In an *in vitro* isolated system that is thrown out of balance by exposure to external stimuli montelukast reduced back IL-10 levels, whereas an asthma patient’s serum is an already unbalanced system brought back closer to balance by montelukast exposure. Our results fall within the first category. Indeed, we found that Montelukast addition decreased IL-10 secretion (Figure ***5***), effectively canceling the accumulative exposure effect.

Altogether, the present efforts lay the basis for future implementation of this setup that combines a recently developed protocol of iPSC-derived bronchial epithelium^45^ and an airway-on-chip model^45^ for the purpose of pollutant toxicity evaluation. Further characterization in the presence of airflow would be of interest to follow up in future work. It has been well established that ALI-based assays and physiologically relevant airflow and directionality offer great advantages over classical cell culture with cells that are submerged underneath culture media at Liquid Liquid Interface (LLI)^57^, such as well-differentiated epithelia with improved ciliary function, mucus secretion and monolayer integrity as well as glycocalyx formation which in turn plays an important role in transmitting shear stress to the epithelial cytoskeleton^18,58,59^, with bidirectional airflow showing more representative muco-ciliary differentiation compared to continuous unidirectional airflow^58^.

## 4. MATERIALS AND METHODS

### 4.1 Model Design and Assembly

We used a previously developed microfluidic device which was assembled as previously described^37^. Briefly, it consists of two Polydimethylsiloxane (PDMS) layers with the top layer consisting of an airway channel in the form and size consistent with adult small airway generations of the bronchial tree. The bottom layer is a basal chamber. The two layers are separated by a porous PET membrane (Corning 408 Falcon® Permeable 0.4 µm Transparent PET Membrane, 353090).

### 4.2 Cell Cultures

#### iPSCs maintenance

iPSCs reprogrammed from healthy donor fibroblasts were kindly gifted by Prof. Lior Gepstein’s laboratory and passaged as previously described^44^. Briefly, cells are cultured in mTeSR™1 media (STEMCELL, Cat#85850) in 6-well culture plates (Nunc Cat#3516) coated with Cultrex RGF BME (R&D Systems Cat#3433-010-01) diluted 1:200 with RPMI-1640 (Sigma Cat#R8758). iPSCs are split routinely when reaching ∼85% confluence. Passaging is performed up to passage 30 provided that expression of the quality control markers (listed below) is maintained and no differentiation is visible upon morphological inspection.

Detachment is achieved by short and robust pipetting in mTeSR following a 7-minute incubation in 1ml/well of EDTA 0.5 mM in PBSX1. Rock inhibitor (Y-27632, R&D systems, Cat#1254/10) at 10 µM is added only during the first 24 hours after passaging. iPSCs quality was routinely verified using RT-PCR (for details see Supporting Data S1) to ensure Yamanaka factors expression (Oct3/4, Sox2, c-Myc, and Klf4)^60^. Routine mycoplasma testing was carried out using isothermal PCR-based mycoplasma detection kit (MycoBlue by Vazyme, Cat#D101).

#### iPSCs to iBCs Differentiation

Differentiation was carried out as previously described^45^ with a few adjustments depicted herein. As to the protocol of epithelial differentiation it noteworthy is the fact that we have introduced a simplified method for cell enrichment in the process of iBCs differentiation and maintenance. Substitution of FACS-sorting for magnetic beads-based columns for enrichment proved to be both time and cost-effective for routine performance. On day 3 after differentiation signal induction successful Definitive Endoderm (DE) was verified by FACS analysis of CXCR4 and CD117 expression (Supplementary ***Figure S1A, Supplementary Table S1***). On day 14 CPM^+^ cells (stained with Mouse Monoclonal Anti-Carboxypeptidase M, Fujifilm, Cat# 014-27501 and Anti-Mouse IgG MicroBeads, Miltenyi Biotec Cat#130-048-402) were selected using either magnetic beads columns (Miltenyi Biotec, Cat# 130-042-401) or BD FACSMelody™ Cell Sorter (BD Biosciences) for comparison. NKX2.1 expression (Supplementary Table S1) correlated with CPM expression and was maintained on day 40 when cells were again enriched for NGFR (Nerve Growth Factor Receptor) and TP63 positive expression (Tumor protein p63). Starting day 14 cells were passaged in 3D as described below (see iBCs maintenance section). Mature iBCs were then cryopreserved (4x10^5^ cells per 1 ml <u>NutriFreez® D10 Cryopreservation Medium, Sartorius, Cat# 05-713-1A).</u>

#### iBCs maintenance

iBCs were maintained as previously described^45^. Briefly, iBCs were routinely grown as 3D organoids within 30 µl domes of Cultrex RGF BME (R&D Systems Cat#3433-010-01) seeded in 24 or 12-well culture dishes and submerged in PneumaCult™-Ex Plus media (STEMCELL, Cat#05040), supplemented with A83-01 (Sigma, Cat# <u>SML0788</u>), DMH1 (Sigma, Cat#D8946) and Y-27632 (R&D systems, Cat#1254/10) at the final concentrations of 1 µM, 1 µM and 10 µM, respectively. Cells were routinely passaged every 10-14 days, up to passage 12. Routine Mycoplasma tests were carried out using isothermal PCR-based mycoplasma detection kit (MycoBlue by Vazyme, Cat#D101).

#### Bronchial epithelia differentiation

Bronchial epithelium was differentiated as previously described^45^ (see ***Table S2, Figure 1b)***). iBCs were seeded in models or trans-wells (24-well, Corning, Cat#CLS3397) at 130,000 cells per model or well, respectively, in iBCs maintenance media for 4 days (in both apical and basal chambers). Media was changed every 2 days. Subsequently cells were transferred into PneumaCult™-ALI Medium (STEMCELL Cat #05001) supplemented with 0.2% Heparin Solution (STEMCELL, Cat# 07980), 0.5% Hydrocortisone Stock Solution (STEMCELL, Cat#07925) and 1% Penicillin/Streptomycin (10,000U/mL) for an additional day (day 4=ALI day 0) and the following day media was removed from the apical chamber to achieve ALI conditions (ALI day 1). By day 8 ciliary beat could clearly be visible microscopically and the culture was used for downstream, experiments no later than ALI day 20.

### 4.3 Epithelial Barrier Integrity

Epithelial Barrier Integrity was evaluated either directly by measuring Trans-Electrical Epithelial Resistance (TEER) in trans-wells or indirectly by measuring apparent permeability (*P*_*app*_) in models. TEER was measured as previously described^61^. Briefly, pre-coated trans-wells (PET membrane surface area of 0.33 cm^2^,6.5 mm diameter and 0.4 um pore size; corning Cat#CLS3397) were seeded with 240,000 iBCs on day 0 of epithelial differentiation. Electrical resistance was measured using chopstick electrodes (MERSSTX01) and the Millicell® ERS-2 Epithelial Volt-Ohm Meter (EMD Millipore). Measurements were performed routinely on days 3-4 following seeding, prior to transferring the culture to ALI conditions. An additional measurement was performed upon epithelial maturation and detection of a visible ciliary beat starting day 11-12 (day 7-8 at ALI). Both electrode poles were fully submerged in growth media. Measurements were carried out at room temperature and three technical repeats were obtained for each trans-well. Tissue TEER (Ohms*cm^2^) was calculated by subtracting the blank resistance (resistance of an uncoated and unseeded well, typically ∼150-300 ohms) from that of the seeded trans-well and multiplying by the surface area of the membrane. A TEER of >400 Ohms*cm^2^ was generally considered sufficient to proceed to ALI conditions.

Apparent Permeability (*Papp*) was measured as previously described^61^ and used to evaluate barrier integrity in models where TEER measurements are more cumbersome. Briefly, the hydrophilic marker sodium fluorescein (Sigma, Cat# F6377) was dissolved in DMSO (stock concentration 1 mg/ml) and introduced into the apical chamber at 50-100 ng/ml in PneumaCult™-ALI Medium. 50 µl were removed from the basal chamber every 30 minutes, with at least six time points collected per assay. Fluorescence was measured in a plate reader in an 384-well microplate, flat bottom, transparent, polystyrol (Greiner Cat#781096)

### 4.4 RT-PCR analysis

Total RNA was extracted using Promega RNA isolation kit (Reliaprep RNA Tissue miniprep, Promega Cat#Z6110). The concentration and quality of RNA were assessed using NanoDrop One spectrophotometer (Thermo Scientific). Subsequently, an RT-PCR was performed using GoTaq® 1-Step RT-qPCR System (Promega, Cat#A6020) where cDNA was generated as part of a single-step real-time amplification reaction combining both Reverse Transcriptase and GoTaq® qPCR master mix containing proprietary fluorescent DNA-binding dye, BRYT Green® Dye. Glyceraldehyde-3-Phosphate Dehydrogenase (GAPDH) was used as the housekeeping gene. Relative gene expression (fold induction) was tested in all cell types compared to that of iPSCs and quantification was performed using the adjusted method of delta-delta CT by Pfaffl, 2001^62^.

### 4.5 Immunohistochemistry and imaging

Both model channels were washed with PBSX1 and subsequently fixed with 4% PFA for 10 minutes. Following fixation, the cells were washed again (PBSX1) and permeabilized using 0.1% Triton X-100, 10 minutes. Alternatively, devices were stored after fixation at 4 °C no longer than 7 days. A 3% BSA blocking buffer was used to block cells at room temperature for 30 minutes, and cells were then stained with primary antibodies in 1% BSA buffer for an additional 60 minutes at room temperature. When needed a secondary antibody in 1% BSA buffer was added to the cells for 10 minutes after three washes in PBSX1 at room temperature. DAPI was used to stain the cellular nuclei after the secondary antibody step, and imaging was performed using spinning disk confocal microscopy (Spinning Disk Confocal Olympus IXplore-Spin). Image processing was performed using ImageJ. A full list of the antibodies used is provided in ***Table S1***.

### 4.6 IL-10 Secretion in Response to Benzene Exposure

IL-10 secretion was assessed using Human IL-10 ELISA Kit (Invitrogen, Cat#KHC0101). Benzene was dissolved at the desired concentration from an 11M stock (Benzene, spectral 99.5%, TCI, Cat#B0020) in growth media after antibiotics removal at least 48 hours prior to the assay. Cells were then exposed by adding the Benzene directly into the apical chamber for 1 hour. Subsequently the apical and basal chambers were washed with fresh media and cells were incubated at ALI for 48 hours before media collection from the basal compartment, or as indicated Where Montelukast addition effect was tested, Montelukast sodium hydrate (Sigma, Cat#SML0101) was diluted in DMSO to a stock concentration of 0.006 mg/ml and added into the basal compartment media at a dilution of 1:1000 as previously described^63^.

### 4.7 Cytotoxicity

#### LDH

To evaluate cytotoxicity by cell membrane integrity, lactate dehydrogenase (LDH) release in the basal medium was measured (LDH-Glo^TM^ Cytotoxicity Assay). Briefly, tested media was collected at a 1:100 dilution into storage buffer as per kit instructions. 50 μl of supernatant and 100 μl freshly prepared LDH detection reagent were added into a 96-well flat-bottomed plate and incubated in the dark for 60 min at room temperature. After adding 50 μL stop solution per well, the luminescence signal was recorded using a microplate reader (Infinite® M200 PRO, Tecan).

#### MTT

Cell metabolic activity as an indicator of viability was measured using MTT Growth Kit (Sigma, Cat#CT02) as per manufacturer instructions. Briefly, 100 µl of media collected from the basal chamber of each trans-well seeded with epithelial cells, treated with Benzene or control media. Following a 4 hour incubation in 37ºC 0.1 mL isopropanol with 0.04 N HCl was added to each well. Within an hour, the absorbance was measured on an ELISA plate reader with a test wavelength of 570 nm and a reference wavelength of 630 nm.

## 5. CONCLUSIONS

This proof-of-concept study demonstrates the potential of an iPSC-derived bronchial epithelium-on-chip model to assess subtle inflammatory effects of VOC exposure and lays the foundation for future risk assessment and improved prevention. We achieve this by designing a BOC-based platform for airborne VOCs effect assessment and show that it is sufficiently sensitive to detect the impact of exposure to even non-toxic VOC concentrations. The pairing of this system with iPSCs-derived bronchial epithelium holds great benefits ranging from reproducibility to improved accessibility and standardization. These results merit further efforts to be invested in the study of how airflow introduction during the process of epithelial differentiation within the model and pollutant exposure would affect the accuracy and quality of data this platform can provide.

We see this as a one of a kind pairing between improved variable isolation and physiological-compatibility resulting in a system can potentially provide a sensitive indication of hazardous outcomes of pollutant exposure that currently cannot be established as such using existing approaches.

## 6. SUPPORTING INFORMATION AVAILABLE

The following files are available free of charge:

Supplementary Tables S1-S3

Supplementary Figures S1-S4

## 7. ACKNOWLEDGMENTS

The authors would like to thank Prof. Lior Gepstein who kindly provided the iPSCs used in this work, and his team members Dr. Irit Huber, Ms. Gil Arbel and Ms. Keren Shavtaiv for their professional guidance and support. The authors also thank Ms. Sivan Yoffe for the assistance with CAD schematics.

## REFERENCES

1. Chronic obstructive pulmonary disease (COPD). https://www.who.int/news-room/fact-sheets/detail/chronic-obstructive-pulmonary-disease-(copd).

2. Ten health issues WHO will tackle this year. https://www.who.int/news-room/spotlight/ten-threats-to-global-health-in-2019.

3. One-third of global population at cancer risk due to elevated volatile organic compounds levels | npj Climate and Atmospheric Science. https://www.nature.com/articles/s41612-024-00598-1.

4. Giuliano, M. et al. Effects of low concentrations of benzene on human lung cells in vitro. Toxicol Lett 188, 130–136 (2009).

5. Koken, P. J. M. et al. Temperature, air pollution, and hospitalization for cardiovascular diseases among elderly people in Denver. Environmental Health Perspectives 111, 1312–1317 (2003).

6. Zhang, X. et al. In vitro biomimetic models for respiratory diseases: progress in lung organoids and lung-on-a-chip. Stem Cell Res Ther 16, 415 (2025).

7. Humbert, M. V. et al. Towards an artificial human lung: modelling organ-like complexity to aid mechanistic understanding. Eur Respir J 60, 2200455 (2022).

8. Frangogiannis, N. G. Why animal model studies are lost in translation. J Cardiovasc Aging 2, 22 (2022).

9. Ro, C. How ‘animal methods bias’ is affecting research careers. Nature (2025) doi:10.1038/d41586-025-00593-3.

10. (PDF) Leveraging Epidemiology to Improve Risk Assessment. ResearchGate (2025) doi:10.2174/1874297101104010003.

11. Artzy-Schnirman, A. et al. Advanced in vitro lung-on-chip platforms for inhalation assays: From prospect to pipeline. Eur J Pharm Biopharm 144, 11–17 (2019).

12. Advanced human-relevant in vitro pulmonary platforms for respiratory therapeutics - ScienceDirect. https://www.sciencedirect.com/science/article/abs/pii/S0169409X21002945.

13. Silva, S., Bicker, J., Falcão, A. & Fortuna, A. Air-liquid interface (ALI) impact on different respiratory cell cultures. European Journal of Pharmaceutics and Biopharmaceutics 184, 62–82 (2023).

14. Direct numerical simulation of particle laden flow in a human airway bifurcation model - ScienceDirect. https://www.sciencedirect.com/science/article/pii/S0142727X16304052.

15. Park, S., Newton, J., Hidjir, T. & Young, E. W. K. Bidirectional airflow in lung airway-on-a-chip with matrix-derived membrane elicits epithelial glycocalyx formation. Lab Chip 23, 3671–3682 (2023).

16. Button, B. & Boucher, R. C. Role of Mechanical Stress in Regulating Airway Surface Hydration and Mucus Clearance Rates. Respir Physiol Neurobiol 163, 189–201 (2008).

17. Loiseau, E. et al. Active mucus–cilia hydrodynamic coupling drives self-organization of human bronchial epithelium. Nat. Phys. 16, 1158–1164 (2020).

18. Plebani, R. et al. Modeling pulmonary cystic fibrosis in a human lung airway-on-a-chip. Journal of Cystic Fibrosis 0, (2021).

19. Sone, N. et al. Multicellular modeling of ciliopathy by combining iPS cells and microfluidic airway-on-a-chip technology. Science Translational Medicine 13, eabb1298 (2021).

20. Benam, K. H. et al. Small airway-on-a-chip enables analysis of human lung inflammation and drug responses in vitro. Nat Methods 13, 151–157 (2016).

21. Johnston, S. L., Goldblatt, D. L., Evans, S. E., Tuvim, M. J. & Dickey, B. F. Airway Epithelial Innate Immunity. Front Physiol 12, 749077 (2021).

22. Oldenburger, M. M. et al. Altered cytokine release of airway epithelial cells in vitro by combinations of respiratory syncytial virus, Streptococcus pneumoniae, Printex 90 and diesel exhaust particles. Environ Res 275, 121392 (2025).

23. de Araújo, A. P., da Costa Rodrigues, T., de Oliveira, M. L. S. & Miyaji, E. N. Cytokine secretion by in vitro cultures of lung epithelial cells, differentiated macrophages and differentiated dendritic cells incubated with pneumococci and pneumococcal extracellular vesicles. Braz J Microbiol 55, 3797–3810 (2024).

24. Mills, P. R., Davies, R. J. & Devalia, J. L. Airway Epithelial Cells, Cytokines, and Pollutants. Am J Respir Crit Care Med 160, S38–S43 (1999).

25. Bonfield, T. L. et al. Normal bronchial epithelial cells constitutively produce the anti-inflammatory cytokine interleukin-10, which is downregulated in cystic fibrosis. Am J Respir Cell Mol Biol 13, 257–261 (1995).

26. Gruzieva, O. et al. Exposure to Traffic-Related Air Pollution and Serum Inflammatory Cytokines in Children. Environ Health Perspect 125, 067007 (2017).

27. Peñaloza, H. F. et al. Opposing roles of IL-10 in acute bacterial infection. Cytokine & Growth Factor Reviews 32, 17–30 (2016).

28. Carlini, V. et al. The multifaceted nature of IL-10: regulation, role in immunological homeostasis and its relevance to cancer, COVID-19 and post-COVID conditions. Front. Immunol. 14, (2023).

29. Lee, C. G. et al. Transgenic Overexpression of Interleukin (IL)-10 in the Lung Causes Mucus Metaplasia, Tissue Inflammation, and Airway Remodeling via IL-13-dependent and - independent Pathways*. Journal of Biological Chemistry 277, 35466–35474 (2002).

30. Saitoh, T. et al. Neutrophil Extracellular Traps Mediate a Host Defense Response to Human Immunodeficiency Virus-1. Cell Host & Microbe 12, 109–116 (2012).

31. Soltys, J., Bonfield, T., Chmiel, J. & Berger, M. Functional IL-10 Deficiency in the Lung of Cystic Fibrosis (cftr−/−) and IL-10 Knockout Mice Causes Increased Expression and Function of B7 Costimulatory Molecules on Alveolar Macrophages1. The Journal of Immunology 168, 1903–1910 (2002).

32. Yoon, H. I. et al. Exposure to volatile organic compounds and loss of pulmonary function in the elderly. European Respiratory Journal 36, 1270–1276 (2010).

33. Bhatnagar, A. Environmental Determinants of Cardiovascular Disease. Circulation Research 121, 162–180 (2017).

34. Son, B., Breysse, P. & Yang, W. Volatile organic compounds concentrations in residential indoor and outdoor and its personal exposure in Korea. Environment International 29, 79–85 (2003).

35. Elias-Kirma, S. et al. In situ-Like Aerosol Inhalation Exposure for Cytotoxicity Assessment Using Airway-on-Chips Platforms. Front Bioeng Biotechnol 8, 91 (2020).

36. Suzuki, S. et al. Differentiation of human pluripotent stem cells into functional airway basal stem cells. STAR Protoc 2, 100683 (2021).

37. Elias-Kirma, S. et al. In situ-Like Aerosol Inhalation Exposure for Cytotoxicity Assessment Using Airway-on-Chips Platforms. Front. Bioeng. Biotechnol. 8, (2020).

38. Nof, E. et al. Human Multi-Compartment Airways-on-Chip Platform for Emulating Respiratory Airborne Transmission: From Nose to Pulmonary Acini. Frontiers in Physiology 13, 1–18 (2022).

39. Raman, T. et al. Quality control in microarray assessment of gene expression in human airway epithelium. BMC Genomics 10, 493 (2009).

40. Gohy, S. et al. Altered generation of ciliated cells in chronic obstructive pulmonary disease. Sci Rep 9, 17963 (2019).

41. Pezzulo, A. A. et al. The air-liquid interface and use of primary cell cultures are important to recapitulate the transcriptional profile of in vivo airway epithelia. Am J Physiol Lung Cell Mol Physiol 300, L25–31 (2011).

42. Renò, V. et al. A Novel Approach for the Automatic Estimation of the Ciliated Cell Beating Frequency. Electronics 9, 1002 (2020).

43. Isogenic induced-pluripotent-stem-cell-derived airway- and alveolus-on-chip models reveal specific innate immune responses | Request PDF. ResearchGate (2025) doi:10.1038/s41551-025-01444-2.

44. Itzhaki, I. et al. Calcium Handling in Human Induced Pluripotent Stem Cell Derived Cardiomyocytes. PLOS ONE 6, e18037 (2011).

45. Suzuki, S. et al. Differentiation of human pluripotent stem cells into functional airway basal stem cells. STAR Protoc 2, 100683 (2021).

46. Current status of IL-10 and regulatory T-cells in cancer - PMC. https://pmc.ncbi.nlm.nih.gov/articles/PMC4322764/.

47. HEALTH EFFECTS. in Toxicological Profile for Benzene (Agency for Toxic Substances and Disease Registry (US), 2007).

48. Dino, P. et al. Release of IL-1β and IL-18 in human primary bronchial epithelial cells exposed to cigarette smoke is independent of NLRP3. Eur J Immunol 54, e2451053 (2024).

49. Guo, X. et al. Benzene metabolites trigger pyroptosis and contribute to haematotoxicity via TET2 directly regulating the Aim2/Casp1 pathway. eBioMedicine 47, 578–589 (2019).

50. Minciullo, P. L., Navarra, M., Calapai, G. & Gangemi, S. Cytokine Network Involvement in Subjects Exposed to Benzene. J Immunol Res 2014, 937987 (2014).

51. Vitucci, E., Cannon, C. l. & Johnson, N. ALI Benzene Exposure Induces Oxidative Stress in the Bronchial Epithelium and Promotes Immune Cell Chemotaxis In Vitro. Am J Respir Crit Care Med 211, A5109–A5109 (2025).

52. Protection from benzene-induced immune dysfunction in mice - ScienceDirect. https://www.sciencedirect.com/science/article/abs/pii/S0300483X22000154.

53. Takanashi, S. et al. Interleukin-10 level in sputum is reduced in bronchial asthma, COPD and in smokers. Eur Respir J 14, 309–314 (1999).

54. Stelmach, I., Majak, P., Jerzynska, J. & Kuna, P. The effect of treatment with montelukast on in vitro interleukin-10 production of mononuclear cells of children with asthma. Clin Exp Allergy 35, 213–220 (2005).

55. Yüksel, B. et al. The effect of treatment with montelukast on levels of serum interleukin-10, eosinophil cationic protein, blood eosinophil counts, and clinical parameters in children with asthma. Turk J Pediatr 51, 460–465 (2009).

56. Lin, Y.-C. et al. Effects of montelukast on M2-related cytokine and chemokine in M2 macrophages. J Microbiol Immunol Infect 51, 18–26 (2018).

57. Baldassi, D., Gabold, B. & Merkel, O. Air-liquid interface cultures of the healthy and diseased human respiratory tract: promises, challenges and future directions. Adv Nanobiomed Res 1, 2000111 (2021).

58. Park, S., Newton, J., Hidjir, T. & Young, E. W. K. Bidirectional airflow in lung airway-on-a-chip with matrix-derived membrane elicits epithelial glycocalyx formation. Lab Chip 23, 3671–3682 (2023).

59. Kesimer, M. et al. Molecular organization of the mucins and glycocalyx underlying mucus transport over mucosal surfaces of the airways. Mucosal Immunol 6, 379–392 (2013).

60. Takahashi, K. & Yamanaka, S. Induction of pluripotent stem cells from mouse embryonic and adult fibroblast cultures by defined factors. Cell 126, 663–676 (2006).

61. Kuehn, A. et al. Human alveolar epithelial cells expressing tight junctions to model the air-blood barrier. ALTEX 33, 251–260 (2016).

62. Pfaffl, M. W. A new mathematical model for relative quantification in real-time RT–PCR. Nucleic Acids Res 29, e45 (2001).

63. Nof, E. et al. Ventilation‐induced epithelial injury drives biological onset of lung trauma in vitro and is mitigated with prophylactic anti‐inflammatory therapeutics. Bioengineering & Transla Med 7, e10271 (2022).

