## Supplementary Data for "iPSCs-derived Bronchial Airways-On-Chip for Assessment of Cytokine Secretion triggered by Volatile Organic Compounds"

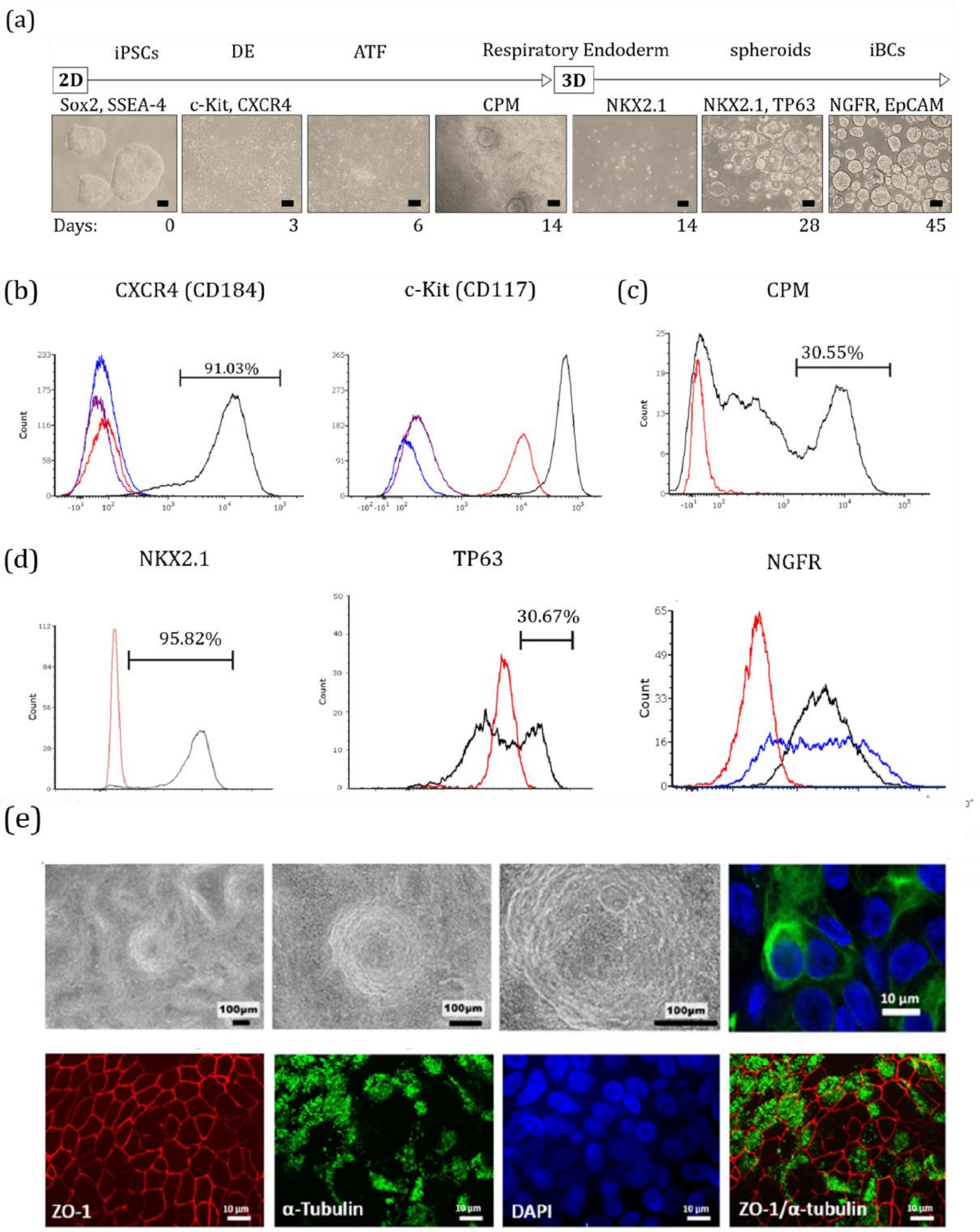

(a)
iPSCs
DE
ATF
Respiratory Endoderm
spheroids
iBCs
2D
3D
Sox2, SSEA-4
c-Kit, CXCR4
CPM
NKX2.1
NKX2.1, TP63
NGFR, EpCAM
Days:
0
3
6
14
14
28
45
(b)
CXCR4 (CD184)
91.03%
c-Kit (CD117)
(c)
CPM
30.55%
(d)
NKX2.1
95.82%
TP63
30.67%
NGFR
Count
(e)
100μm
10 μm
ZO-1
α-Tubulin
DAPI
ZO-1/α-tubulin

**Figure S1: iBCs detailed differentiation stages, sorting strategy and**
**(a)** Summary of the iPSCs to iBCs differentiation process. Scale bars of 100 µm ad stage markers are indicated at the relevant stages. **(b)** Expression of DE Markers CD184 (CXCR4) and CD117 (c-kit) on differentiation day 3 following DE signal induction. DE (—) and iPSCs (—) were stained with primary antibodies (left panel CXCR4, right panel c-kit) conjugated to APC or PE fluorophores respectively. Unstained DE (—) and iPSCs (—) serve as respective controls. Representative results of FACS analysis are shown (n=2). **(c)** Day 14 of differentiation cells were tested for CPM-expression (—) with iPSCs as a negative control (—). Live cells were stained with the cell permeant dye Calcein blue AM dye (not shown). CPM-positive, live cells were enriched and comprised 33.28% of the population. Representative result of two independent repeats (n=2). **(d)** Day 40 onward, representative analysis of routine analysis. NKX2.1 (left panel; iPSCs —, iBCs —), TP63 (center panel; iPSCs —, iBCs —) and NGFR (left panel) expression is monitored during routine passaging and NGFR-positive cells are enriched from 69.26% (—) to 90.75% (—). Secondary antibody only control (—) for background intensity and enrichment percentage gating. **(e) Top panel:** Muco-ciliary patterning indicating active mucus production and ciliary beat in inserts. Starting day 8 of ALI a ciliary beat can be visualized in brightfield. Mucus production precedes it and muco-ciliary patterning takes on the characteristic swirl form. Mucus production is quantified by MUC5AC staining of goblet cells stained green (right). **Bottom panel:** Mature Bronchial Epithelium Morphology. ZO-1 as a tight junction marker (red; left) together with α-tubulin staining of the cilia are characteristic features of a healthy bronchial epithelium.

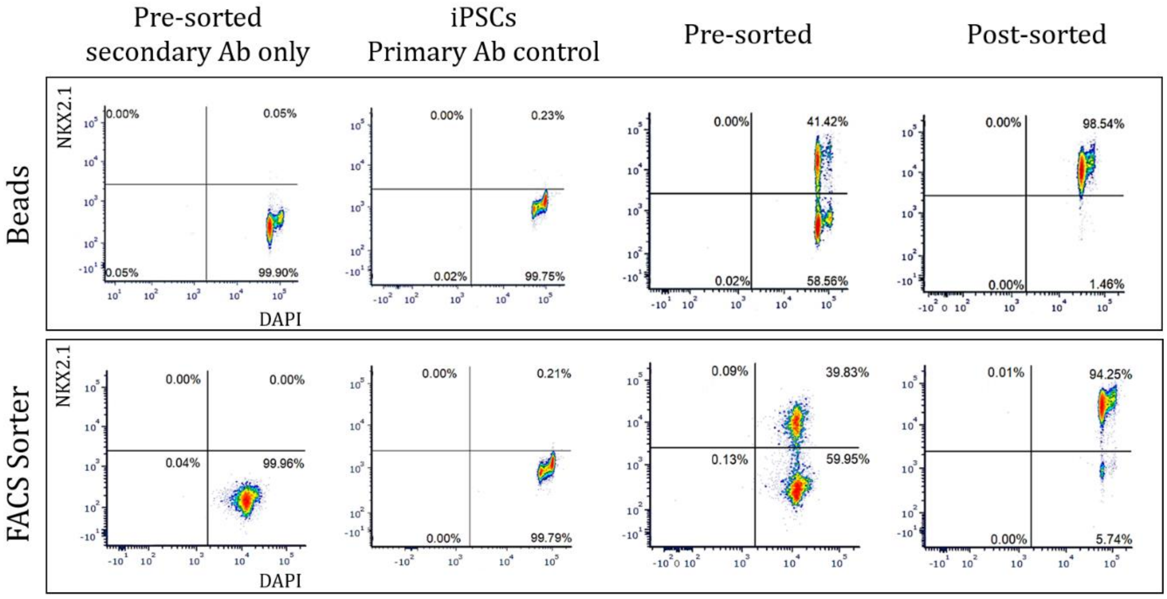


**Figure S2: FACS-sorting versus magnetic beads enrichment of CPM-positive cells expressing**

Comparison between FACS sorting and magnetic beads to isolate NKX2.1-positive cells on days 15 and 4 of differentiation, respectively. Separation was performed based on CPM membranal marker expression in live cells. Following separation cells were fixed to allow for NKX2.1 nuclear transcription factor staining and evaluation. Beads methods had achieved 98.54% NKX2.1-positive cells, while FACS sorter achieved an enrichment of 94.25%.

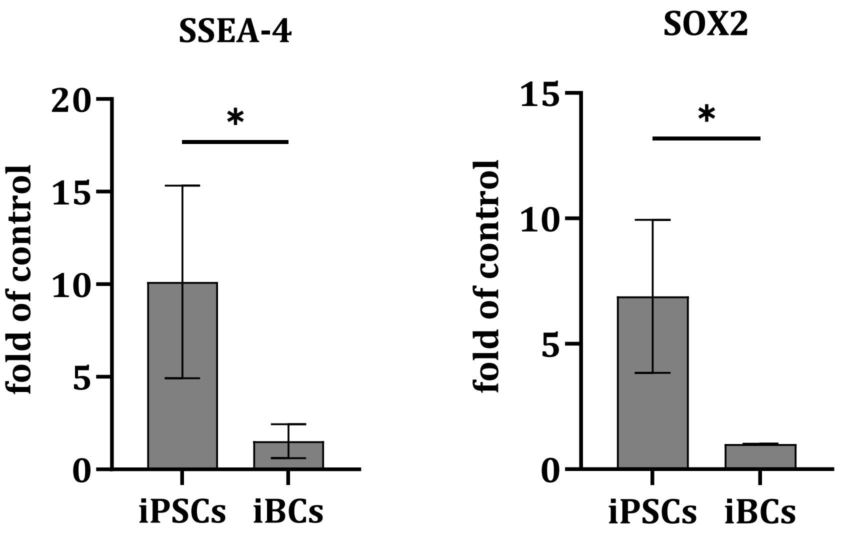


**Figure S3: iPSCs pluripotency markers expression validation**
Pluripotency markers SOX2 and SSEA-4 expression is increased in iPSCs compared to that of iBCs. Unpaired t-test was significant (*p-value<0.05, n=3). Data is normalized to a single repeat of iPSCs.

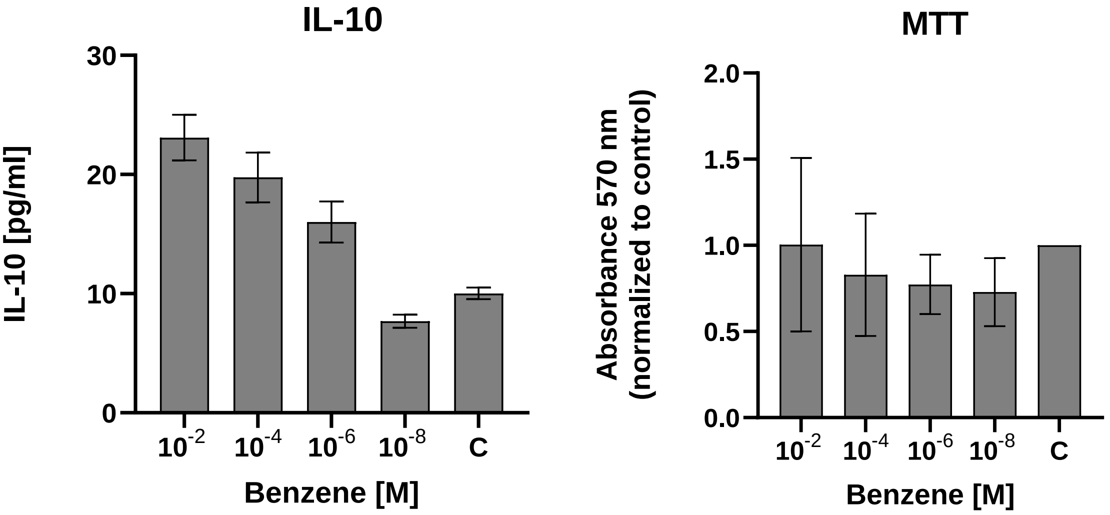


**Figure S4: IL-10 secretion and viability assay (MTT) of iPSCs-derived bronchial epithelia upon Benzene exposure in trans-wells**

Table S1. Antibodies list

| **Target** | **Antibody** | **Source** | **Identifier** |
|---|---|---|---|
| CD117/c-kit | PE Anti-c-Kit antibody [104D2] | Abcam | ab111245 |
| CXCR4 | APC Anti-CXCR4 antibody [12G5] | Abcam | ab270635 |
| CPM | Mouse Monoclonal Anti-CPM (Carboxypeptidase M) | Fujifilm | 014-27501 |
| Anti-Mouse IgG MicroBeads | Anti-Mouse IgG Micro Beads for magnetic column | Miltenyi Biotec | 130-048-402 |
| TTF1/NKX2.1 | Rabbit Anti-TTF1/NKX2-1 antibody [SP141] | Abcam | ab227652 |
| NGFR/CD271 | Mouse anti-human CD271 (NGFR) | BioLegend | BLG-345102 |
| TP63 | p63 Recombinant Rabbit Monoclonal Antibody | Abcam | ET7110-47 |
| EpCAM | CD326 MicroBeads, human | Miltenyi Biotec | 130-061-101 |
| Anti-Rabbit IgG | Alexa Fluor® 488 AffiniPure™ Goat Anti-Rabbit IgG | Jackson Immunostaining | 111-545-144 |
| Anti-Rabbit IgG | Alexa Fluor® 647 AffiniPure™ Goat Anti-Rabbit IgG | Jackson Immunostaining | 111-605-144 |
| Anti-Mouse IgG | Alexa Fluor® 647 AffiniPure™ Goat Anti-Mouse IgG | Jackson Immunostaining | 115-605-062 |
| Anti-Mouse IgG | Alexa Fluor® 488 AffiniPure™ Goat Anti-Mouse IgG | Jackson Immunostaining | 115-545-062 |

Table S2. Time course of iBCs to Mature Epithelium differentiation at ALI

| **DAY 0** | **DAY1-DAY3** | **(TEER>400)** | **ALI** | **ALI** | **DAY7-8 ALI** | **DAY17 ALI** |
|---|---|---|---|---|---|---|
| **Seeding** | **Feeding** | **ALI media** | **Full ALI** | **Feeding** | **Feeding** | **Collection** |
| iBCs media in both chambers | Change both chambers every other day | ALI media in both chambers | ALI media in basal chamber ONLY! | Change media in basal chamber every other day | Change media in basal chamber every other day<br>Cilliary beat becomes visible at X400 mag under light mic | ELISA/Staining/RNA collection |

Table S3. Primer Sequences used for RT-PCR

| Primer | Sequence |
|---|---|
| GAPDH-FW | ACCTGACCTGCCGTCTAGAA |
| GAPDH-RV | TCCACCACCCTGTTGCTGTA |
| **iBCs** | |
| NKX2.1-FW | CGGCATGAACATGAGCGGCAT |
| NKX2.1-RV | GCCGACAGGTACTTCTGTTGCTTG |
| TP63-FW | CCCTCCAACACCGACTACCC |
| TP63-RV | CACCGCTTCACCACCTCCGT |
| NGFR-FW | CTTCTGGGGGTGTCCCTT |
| NGFR-RV | ACTCACCGCTGTGTGTGTA |
| EpCAM-FW | CGCAGCTCAGGAAGAATGTG |
| EpCAM-RV | TGAAGTACACTGGCATTGACG |
| **Epithelium** | |
| KRT5-FW | TCCAGTGTGTCCTTCCGAAGT |
| KRT5-RV | TGCCTCCGCCAGAACTGTA |
| ZO-1-FW | TGCCATTACACGGTCCTCTG |
| ZO-1-RV | GGTTCTGCCTCATCATTTCCTC |
| Occludin-FW | TGCATGTTCGACCAATGC |
| Occludin-RV | AAGCCACTTCCTCCATAAGG |
| TUBA1A-FW | CCACAGTCATTGATGAAGTTCG |
| TUBA1A-RV | GCTGTGGAAAACCAAGAAGC |
| **iPSCs** | |
| SSEA-4-FW | TGGACGGGCACAACTTCATC |
| SSEA-4-RV | GGGCAGGTTCTTGGCACTCT |
| c-MYC-FW | GCGTCCTGGGAAGGGAGATCCGGAGC |
| c-MYC-RV | TTGAGGGGCATCGTCGCGGGAGGCTG |
| OCT3/4-FW | CTTGCTGCAGAAGTGGGTGGAGGAA |
| OCT3/4-RV | CTGCAGTGTGGGTTTCGGGCA |
| SOX2-FW | GCACATGAAGGAGCACCCGGATTA |
| SOX2-RV | CGGGCAGCGTGTACTTATCCTTCTT |
| KLF4-FW | ACGATCGTGGCCCCGGAAAAGGACC |
| KLF4-RV | TGATTGTAGTGCTTTCTGGCTGGGCTCC |
